# Temperature and species-dependent regulation of browning in retrobulbar fat

**DOI:** 10.1101/2020.10.12.333807

**Authors:** Fatemeh Rajaii, Dong Won Kim, Jianbo Pan, Nicholas R. Mahoney, Charles G. Eberhart, Jiang Qian, Seth Blackshaw

**Affiliations:** Department of Ophthalmology, Wilmer Eye Institute, Johns Hopkins University School of Medicine, Baltimore, MD; Solomon H. Snyder Department of Neuroscience, Kavli Neuroscience Discovery Institute, Johns Hopkins University School of Medicine, Baltimore, MD; Department of Pathology, Johns Hopkins University School of Medicine, Baltimore, MD; Department of Neurology, Johns Hopkins University School of Medicine, Baltimore, MD; Institute for Cell Engineering, Johns Hopkins University School of Medicine, Baltimore, MD; Kavli Neuroscience Discovery Institute, Johns Hopkins University School of Medicine, Baltimore, MD

## Abstract

Retrobulbar fat deposits surround the posterior retina and optic nerve head, but their function and origin are obscure. We report that mouse retrobulbar fat is a neural crest-derived tissue histologically and transcriptionally resembles interscapular brown fat. In contrast, human retrobulbar fat closely resembles white adipose tissue. Retrobulbar fat is also brown in other rodents, which are typically housed at temperatures below thermoneutrality, but is white in larger animals. We show that retrobulbar fat in mice housed at thermoneutral temperature show reduced expression of the brown fat marker *Ucp1*, and histological properties intermediate between white and brown fat. We conclude that retrobulbar fat can potentially serve as a site of active thermogenesis, that this capability is both temperature and species-dependent, and that this may facilitate regulation of intraocular temperature.

## Introduction

Retrobulbar fat surrounds the posterior pole of the eye and optic nerve head in all vertebrates studied to date. While adipose tissues in the trunk have been extensively studied, the function and molecular properties of retrobulbar fat are much less well understood, despite the fact that increased volume of retrobulbar fat is a major complication in Graves disease^1^. Retrobulbar fat in cats is less metabolically active than other adipose tissues^2^, and is thus not thought to contribute to energy storage and homeostasis. Instead, retrobulbar fat has been proposed to act as both thermal and mechanical insulation, preventing the loss of heat that is needed to sustain retinal and optic nerve function^3–5^ and supporting the globe and other orbital tissues during eye movements^6^ and trauma^7–9^.

In addition to its role in energy storage, adipose tissue also plays a central role in regulating body temperature^10^. Recent studies have raised the possibility that orbital fat might also engage in active thermogenesis, although any such function is likely to be species-dependent. White adipose tissue, which comprises the bulk of body fat and primarily serves as a site of energy storage, also acts passively as an insulator to reduce heat loss. Brown adipose tissue, in contrast, undergoes active thermogenesis in response to cold. In mice, retrobulbar fat has been reported to express the brown fat marker *Ucp1,* and to upregulate *Ucp1* expression in a model of Grave’s Disease induced by injection of antibodies to thyrotropin receptor (TSHR)^11, 12^. In hibernating 13-lined ground squirrels, however, retrobulbar fat was reported to histologically resemble brown fat but not express *Ucp1*^13^. Finally, human retrobulbar fat in both healthy controls and thyroid eye disease patients morphologically resembles white fat^14, 15^.

While this potentially suggests a thermogenic function for retrobulbar fat in mice, this has not been demonstrated, nor have the reasons behind these potential species-specific differences been explored. More generally, several lines of evidence suggest that retrobulbar fat may be intrinsically different from trunk adipose tissue, most notably in the fact that it is unaffected by genetic diseases such as lipodystrophy that compromise the viability or function of better-studied adipose tissues^16^. The extent to which retrobulbar fat resembles other brown and white fat depots at the molecular level remains unknown for both mice and humans. These intrinsic differences may result from differences in the developmental origin of retrobulbar fat relative to trunk adipose tissues. However, although many other periorbital tissues -- such as cranial bones, ocular muscles, and periocular mesenchyme -- are known to be derived from embryonic neural crest^17–19^, the developmental origin of retrobulbar fat has not been characterized.

In this study, we set out to define the developmental origins of murine retrobulbar fat and transcriptionally profile retrobulbar fat in mice and humans. We find that retrobulbar fat is a neural crest-derived tissue that expresses a small subset of genes that are not detected in trunk fat deposits. In mice, retrobulbar fat resembles interscapular brown fat both histologically and transcriptionally closely, while in humans, retrobulbar fat resembles white adipose tissue. Histological analysis using a range of mammalian species revealed that retrobulbar fat is brown in rodents, which are typically housed at temperatures below thermoneutrality, but is white in larger animals. We further show that in mice housed at thermoneutral temperature retrobulbar fat shows reduced expression of brown fat markers and histological properties that are intermediate between white and brown fat. We conclude that retrobulbar fat can potentially serve as a site of active thermogenesis in rodents, and that this may help maintain intraocular temperature at levels sufficient for proper visual function.

## Results

### Mouse orbital fat is derived from embryonic neural crest

We first used genetic fate mapping to determine whether retrobulbar fat arises from neural crest-derived progenitors, using the neural crest-specific *Wnt1-Cre* transgene^18^ in combination with the Cre-dependent *CAG-lox-stop-lox-Sun1-GFP* transgene^20^ to irreversibly label neural crest-derived cells. At 9 weeks of age, we observed that GFP expression selectively marked the nuclear envelope of retrobulbar adipocytes in *Wnt1-Cre;CAG-lox-stop-lox-Sun1-GFP* mice but not in Cre-negative *CAG-lox-stop-lox-Sun1-GFP* littermate controls (Figure 1A, 1B), demonstrating that murine orbital fat is derived from *Wnt1*-expressing neural crest progenitors.

**Figure 1.**
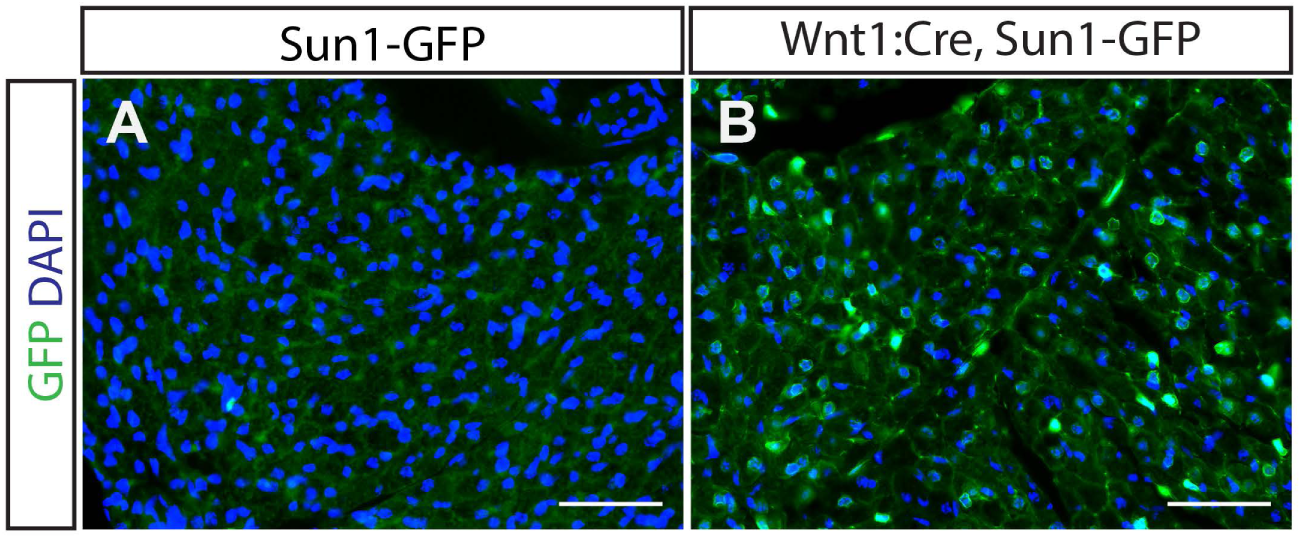
Murine retrobulbar orbital fat derives from Wnt1-positive precursors. **A.** Immunohistochemistry of cryosection of orbital soft tissue from a Cre-negative, Sun1-GFP mouse reveals absent GFP staining of orbital fat. **B.** Immunohistochemistry of cryosection of orbital soft tissue from *Wnt1:Cre;Sun1-GFP* mice reveals presence of nuclear GFP signal in orbital fat. Scale bars = 50 μm.

### The mouse orbital fat transcriptome closely resembles interscapular brown fat

In order to better understand the function of orbital fat compared to other fat depots, we performed bulk RNA sequencing (RNA-Seq) to compare the transcriptomes of mouse orbital fat, interscapular brown fat, and two white fat depots (epididymal fat and inguinal fat) from 9-10 week old CD1 male mice. Principal component analysis demonstrates that orbital fat clusters more closely to brown fat than to white fat depots such as epididymal fat and inguinal fat, which in turn both cluster closely to one another (Figure 2A). Sample-wise correlation likewise shows that orbital fat is more closely related to brown fat than to either epididymal fat or inguinal fat, which are in turn closely related to each other (Figure 2B). Finally, heat map analysis of gene expression also reveals extensive overlap between epididymal fat or inguinal fat, but minimal overlap between those depots and either brown fat or orbital fat, while brown fat and orbital fat in turn show similar patterns of gene expression (Figure 2C).

**Figure 2.**
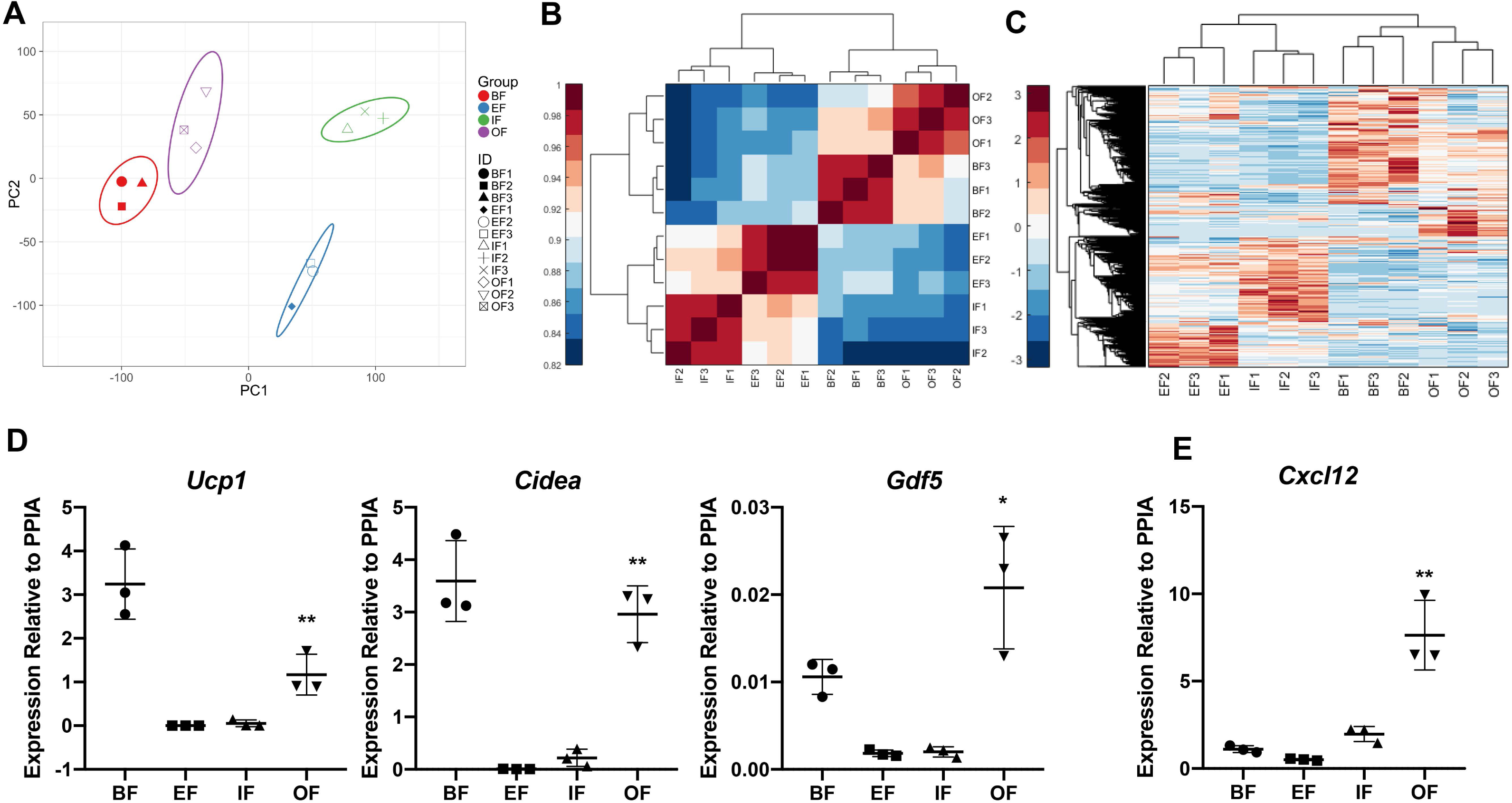
Transcriptomic analysis of murine fat depots reveals significant similarity between orbital fat and interscapular brown fat. **A.** Principal component analysis of transcriptome data from mouse brown fat (BF), epididymal fat (EF), inguinal fat (IF) and orbital fat (OF) demonstrates that BF and OF segregate more closely to each other than to EF and IF. **B.** Sample-wise correlation of transcriptome data from BF, EF, IF and OF demonstrates that transcriptome of OF is more closely correlated with that of BF than with the transcriptomes of EF and IF. **C.** Heat map of transcriptome data from mouse BF, EF, IF and OF demonstrates more significant shared gene expression between OF and BF. There is less significant overlap between OF and either EF or IF. **D.** qPCR validates transcriptome data, demonstrating that *Cxcl12* transcript variant 2 is enriched in orbital fat (OF) compared to brown fat (BF), epididymal fat (EF), and inguinal fat (IF). ** indicates p<0.0001 by one-way Anova. **E.** qPCR demonstrates that brown fat markers *Ucp1*, *Cidea*, and *Gdf5* are enriched in murine BF and OF compared to EF and IF. ** indicates p<0.0001 by one-way Anova. * indicates p=0.006 by one-way Anova.

To further confirm that retrobulbar fat also expressed genes enriched in brown fat, we also performed qRT-PCR analysis for the brown fat markers *Ucp1*, *Cidea*, and *Gdf5,* and confirmed these genes were indeed strongly enriched in orbital fat relative to white fat, and were expressed at similar levels to those seen in brown fat (Figure 2D).

### Identification of genes selectively expressed in orbital fat

A small number of genes are specifically enriched in orbital fat and not detected in other adipose tissues (Table S1). Since mouse orbital fat constitutes a small portion of the orbital volume, and is enveloped by extraocular muscles which may potentially contaminate dissection, RNA-Seq analysis of extraocular muscles was also performed (Figure S1). We found that the extraocular muscle transcriptome did not cluster near that of orbital fat, and that putative retrobulbar fat-enriched transcripts did not arise as the result of dissection contamination.

In order to identify genes that are specific to orbital fat, expression levels between the four fat depots and extraocular muscle (FDR<0.1, fold change>2) were compared. Nine genes were found to be significantly upregulated in orbital fat compared to the other tissues (Table S1). Two of these -- *Cxcl12* and *Slc16a2* -- are particularly noteworthy because of potential links to the pathogenesis of Graves disease, in which inflammation and adipogenesis are selectively observed in retrobulbar fat, but not other adipose tissues. *Cxcl12* encodes a chemokine that can bind two different receptors, CXCR4 and ACKR3, activating pathways that promote inflammation and that regulates cell proliferation, survival, and migration during development^21^. *Cxcl12* and its receptors are dysregulated in a range of autoimmune disorders^22^. This analysis identified several genes, including the cytokine *Cxcl12* that were selectively enriched in retrobulbar fat relative to other white and brown fat depots. *Slc16a2* encodes MCT8, a high-affinity thyroid hormone transporter^23^. We found that *Slc16a2* is expressed in brown, inguinal and epididymal fat depots as well as extraocular muscle, however expression levels are significantly higher in orbital fat (Tabel S1). We used qPCR to confirm selective enrichment *Cxcl12* in retrobulbar fat compared to brown, epididymal and inguinal fats (Figure 2E).

Two other retrobulbar fat-enriched genes that we identified have interesting disease associations, *Gldn* and *Serpini1*. *Gldn* is expressed in nodes of Ranvier and pathologic variants of which cause lethal arthrogryposis multiplex congenita^24^. *Serpini1* is a serine protease inhibitor primarily expressed in the central nervous system, pathologic variants of which are associated with progressive myoclonus epilepsy^25^. *Serpini1* has also been described as a tumor suppressor in gastric tumors as well as having a putative role in epithelial-mesenchymal transition in colorectal cancer models^26, 27^. The functions of *Gldn* and *Serpini1* in orbital fat are unknown. Gene loci with unknown functions were also identified. In addition, seventeen genes were demonstrated to be significantly down-regulated in orbital fat compared to the other adipose tissues (Table S1).

### Retrobulbar fat is brown in rodents but not in larger mammals

Brown fat is a specialized thermogenic tissue that is present in only a few defined regions in the trunk^28^, and these findings raised the question of what function brown fat might play in the retrobulbar region. One possibility is that adaptive thermogenesis might serve to maintain the posterior retina and optic nerve head at consistently high temperatures to maintain consistent conditions for neurotransmission, and thus visual function, in the face of low environmental temperatures. Since small mammals such as mice are typically housed below thermoneutral temperatures, where adaptive thermogenesis must take place to maintain normal body temperature, we hypothesized that orbital fat might also be brown in other rodents, but be white in larger animals, whose large eyes make their retinas and optic nerve heads more resistant to the effects of low ambient temperatures.

To test this, we examined hematoxylin and eosin (H&E) stained sections of orbital fat obtained from adult hamsters, cats, dogs, sheep, and goats. We observe that orbital adipose tissue of mouse and hamster is brown adipose tissue, while orbital adipose tissue in cat, dog, sheep, and goat are white adipose tissue (Figure 3).

**Figure 3.**
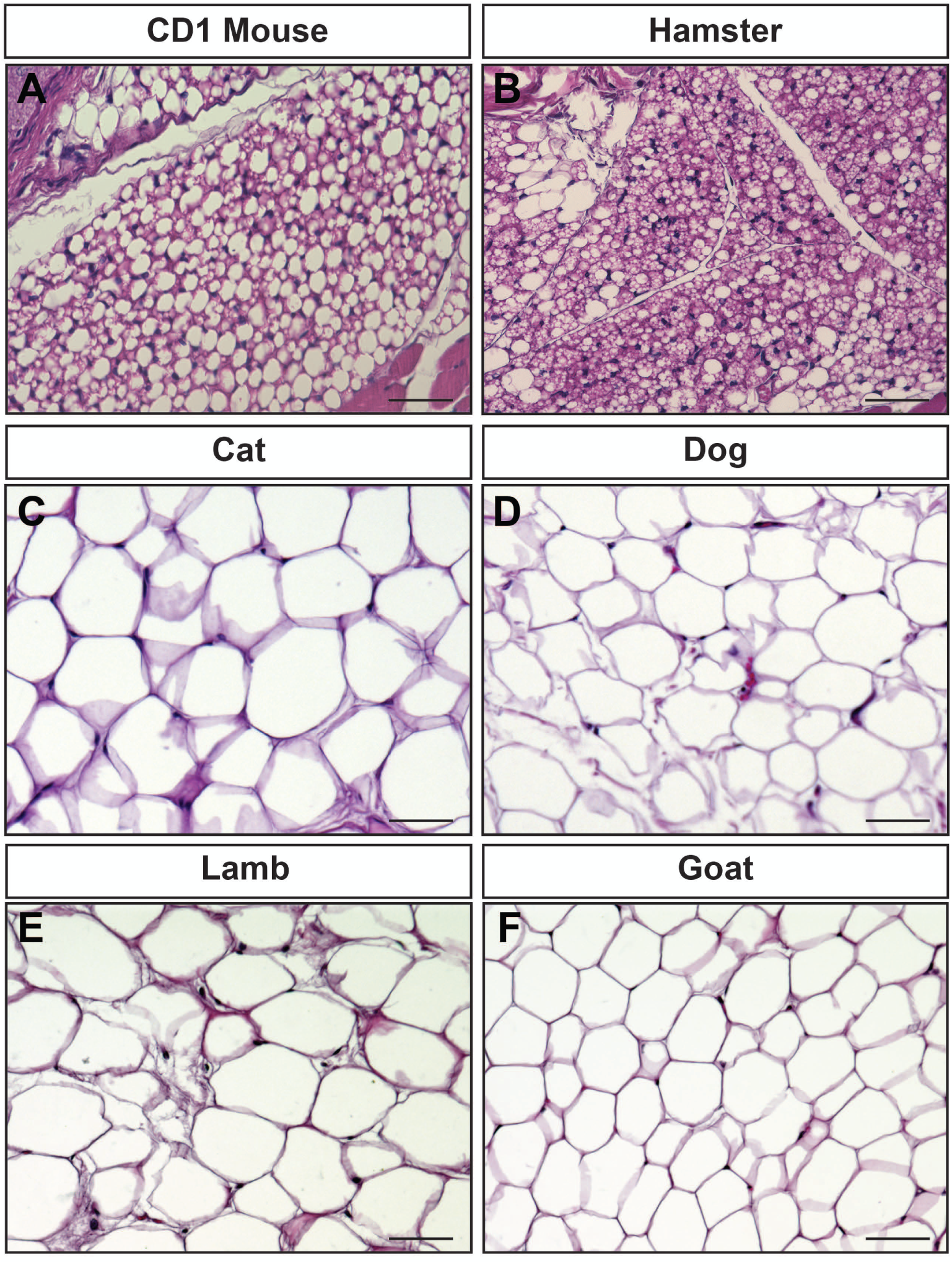
Histologic analysis of orbital fat from different species reveals that brown fat is present in rodent species, but not the other mammals examined. **A.** H&E stained section of CD1 mouse orbital fat demonstrates a mixture of cells with small lipid vacuoles interspersed with cells with large, single lipid vacuoles. **B.** H&E stained section of hamster orbital fat reveals the presence of adipose cells containing multiple small lipid vacuoles. **C.** H&E stained section of cat orbital adipose tissue reveals cells with large lipid vacuoles. **D.** H&E stained section of dog orbital fat reveals cells with large lipid vacuoles. **E.** H&E stained section of lamb orbital fat reveals cells with large lipid vacuoles. **F.** H&E stained section of goat orbital adipose tissue reveals cells with large lipid vacuoles. Scale bars = 50 μm.

Thermoneutral temperatures for mouse (30°C) and hamster (25-32°C) are well above the 22°C degree temperature at which they were housed^29, 30^. These results thus closely match our prediction, where small animals housed at temperatures below thermoneutral exhibiting brown retrobulbar fat, and larger animals exhibiting white retrobulbar fat.

### Murine orbital adipose tissue undergoes whitening in animals housed at thermoneutral temperatures

In order to determine whether murine orbital adipose tissue is intrinsically brown or whether this property is regulated by environmental temperature, mice were housed at thermoneutral temperature for 4 weeks prior to analysis of orbital and interscapular adipose tissue by H&E staining. Examination of histologic sections of tissue reveals that both the interscapular brown adipose tissue as well as the orbital adipose tissue undergo substantial whitening after thermoneutral adaptation, though some cellular heterogeneity was observed, with histologic features of brown fat retained in a subset of cells (Figure 4A-C). SmfISH demonstrated that *Ucp1* expression was substantially reduced in the great majority of cells interscapular brown fat under thermoneutral conditions, as previously reported (Figure 4D-I)^31^. Retrobulbar adipocytes likewise showed a reduction in *Ucp1* expression (Figure J-O), consistent with histological data and also showed a reduction in *Gdf5* (Figure 4P-U). A strong reduction in *Cxcl12* expression was also noted at thermoneutrality, demonstrating that the expression of genes specifically enriched in retrobulbar adipose tissue, compared to other fat depots, may also be temperature-regulated (Figure 4P-U).

**Figure 4.**
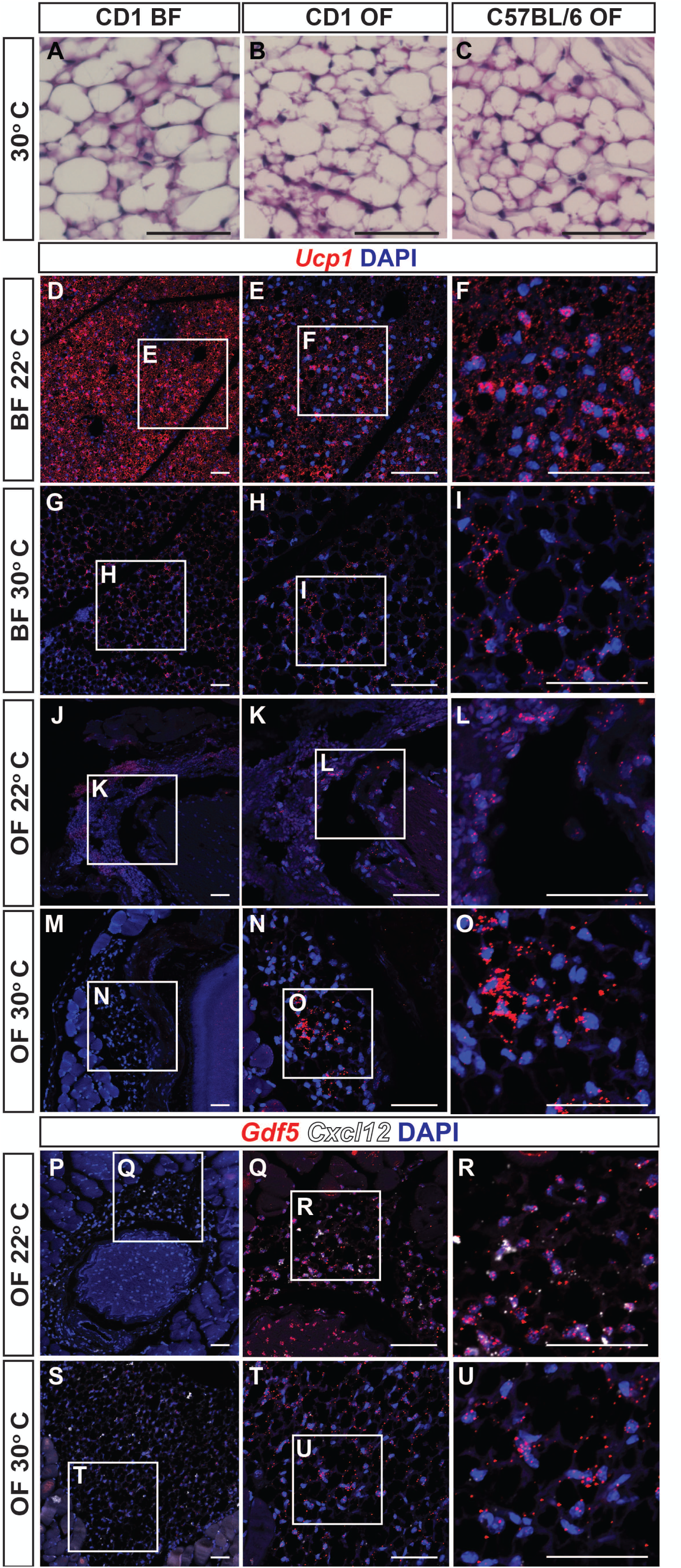
Mouse orbital fat resembles interscapular brown fat after thermoneutral adaptation. **A.** H&E section of interscapular brown fat (BF) from CD1 mice housed at 30°C for 4 weeks demonstrates increased proportion of adipocytes with large lipid droplets characteristic of white adipose tissue. **B.** H&E section of orbital fat (OF) from CD1 mouse housed at 30°C for 4 weeks demonstrates increased proportion of adipocytes with large lipid droplets characteristic of white adipose tissue. **C.** H&E section of OF from C57BL/6 mouse housed at 30°C for 4 weeks demonstrates increased proportion of adipocytes with large lipid droplets characteristic of white adipose tissue. **D-I.** smfISH demonstrates decreased *Ucp1* expression in interscapular brown fat of mice adapted to thermoneutral temperatures (30°C) compared to those housed near room temperature (22°C). **E.** Higher magnification of D. **F.** Higher magnification of E. **H.** Higher magnification of G. **I.** Higher magnification of H. **J-O.** smfISH demonstrates decreased *Ucp1* expression in orbital fat of mice adapted to thermoneutral temperatures (30°C) compared to those housed near room temperature (22°C). **K.** Higher magnification of J. **L.** Higher magnification of K. **N.** Higher magnification of M. **O.** Higher magnification of N. **P-U.** SmfISH demonstrates decreased *Gdf5* and *Cxcl12* expression in orbital fat of mice adapted to thermoneutral temperatures (30°C) compared to those housed near room temperature (22°C). **Q.** Higher magnification of P. **R.** Higher magnification of Q. **T.** Higher magnification of S. **U.** Higher magnification of T. Scale bars = 50 μm.

### Human orbital fat is white, but has an expression profile that is distinct from trunk white adipose tissue

Consistent with previous reports and observations in other large mammals, we observe that retrobulbar orbital adipose tissue from healthy human donors resembles white fat, consisting of adipocytes containing a single large lipid droplet (Figure 5A)^14, 15^. Analysis of RNA-Seq data of prolapsed retrobulbar fat obtained from healthy donors demonstrated that brown fat markers such as *UCP1*, *ZIC1*, and *DIO2* were expressed at levels comparable to that seen in trunk white adipose tissue^32^. Overall, however, human orbital fat has a distinct transcriptional profile, expressing previously identified markers of both white and beige fat (Figure 5B)^33–36^, while also expressing orbital fat-enriched markers identified in mouse (Figure 2). Interestingly, treatment of cultured human orbital fat stem cells derived from both thyroid eye disease patients and healthy controls with bimatoprost, a prostanoid drug associated with prostaglandin-associated periorbitopathy in humans, increases mitochondrial staining and induces expression of *UCP1*, indicating a possible browning of orbital adipocytes which may contribute to the phenotype of periocular fat atrophy^37^. Consistent with its expression of a subset of beige adipose markers, human orbital fat thus also displays some capacity for browning *in vitro*.

**Figure 5.**
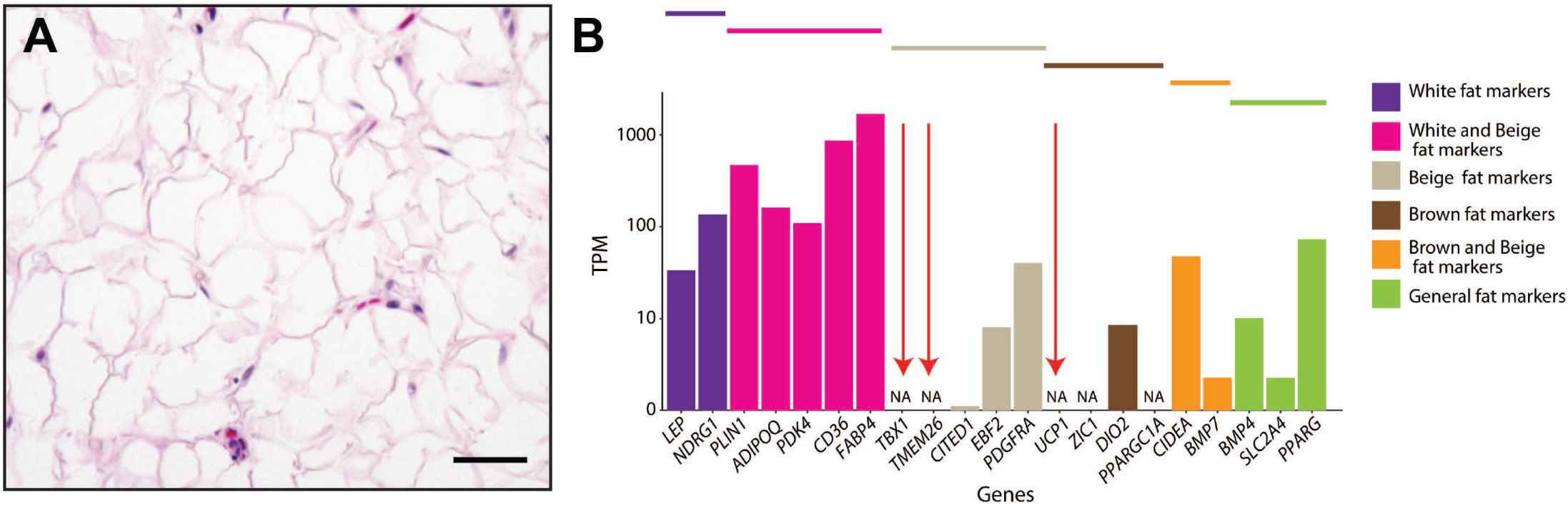
Human orbital adipose tissue is histologically similar to white fat, but expresses white, beige, and brown fat markers. **A.** H&E stained section of human orbital fat reveals cells with large lipid vacuoles. Scale bar = 50 μm. **B.** Comparison of transcriptome data from human orbital fat to publicly available data from various human samples.

## Discussion

In this study, we provide the first complete transcriptomic profiling of murine orbital fat and compare it to other adipose depots to better characterize the tissue, concluding that murine orbital fat most closely resembles brown fat. In addition, we provide evidence that orbital adipose tissue in mammals is derived from neural crest precursors, and provide evidence that neural crest is able to give rise to brown adipose tissue. We have performed comparative anatomy to establish that the presence of brown fat in the orbit is not conserved among mammalian species including cat, dog, goat, sheep, and human orbital fat. We further demonstrate that at thermoneutral temperatures, orbital fat maintains the histologic signature of brown fat, but contains an increased number of white adipocytes.

Though other groups have demonstrated the expression of *Ucp1* in mouse orbital adipose tissue, mouse orbital fat has been referred to as both brown and beige fat in the literature^12, 38, 39^, with little discussion of the distinction or its functional implications. Brown adipose tissue functions in thermogenesis, and in maintaining core body temperature under low ambient temperature. It is likely that the presence of highly specialized brown fat in the rodent orbit has functional significance. Bony fish and sharks which exhibit cranial endothermy -- the ability to maintain elevated brain and eye temperatures -- accomplish this using adaptive thermogenesis from specialized muscle cells or a counter-current heat exchanger to maintain elevated temperatures^3–5, 40, 41^.

These specialized tissues are similar to brown fat in mammals in that they have ample blood supply, high mitochondrial content and high content of cytochrome c^3^.

Increased retinal temperatures allows for faster rates of neurotransmission and potentially improved function^42^, and *in vitro* electrophysiology of the swordfish retina has demonstrated improved temporal resolution as measured by the flicker fusion frequency^43^. Mice housed under standard conditions at 22°C must expend nearly a third of their energy in maintaining core body temperature, and lower temperatures in the retina and optic nerve result in substantially reduced processing speed and visual acuity^42, 44^. We hypothesize that brown fat in the rodent orbit serves a similar function to the specialized muscle or countercurrent heat exchanger in some fish, and may be important for allowing optimal visual function under low external temperatures.

## Methods

### Fate mapping of murine orbital fat

The use of animals for these studies, and all relevant procedures used, were performed in accordance with regulations and guidelines approved by the Johns Hopkins Animal Care and Use Committee. We crossed Wnt1:Cre (JAX stock #003829, Jackson Labs) mice with Sun1-GFP (Jax Stock #021039, Jackson Labs) reporter mice to allow fate mapping of neural crest-derived cells in orbit. Six to 9 week old males were perfused with 4% paraformaldehyde (PFA), decapitated, and the scalp degloved. The skulls were soaked in 4% PFA overnight, washed, and decalcified in 10% EDTA overnight prior to embedding in VWR Clear Frozen Section Compound (VWR, West Chester, PA). Blocks were sectioned at 16 um at stained using rabbit-anti GFP Polyclonal Antibody (Thermo Fisher Scientific, catalog # A-6455, RRID AB_221570) with Goat anti-Rabbit IgG (H+L) Highly Cross-Adsorbed Secondary Antibody, Alexa Fluor 488 (Thermo Fisher Scientific, catalog # A-11034, RRID AB_2576217).

### Transcriptome Analysis

Nine-week old CD-1 mice were euthanized and their OF, inguinal, epididymal, and brown fat isolated. RNA was extracted from each fat depot using Trizol and RNeasy mini kits (Qiagen, Hilden, Germany). Fat from 5 mice was pooled into each sample. A bioanalyzer was used to perform quality control. cDNA libraries were prepared for samples with RIN>7 and RNA sequencing performed using Illumina Nextseq500 on three samples per fat depot. The RNA-seq reads were mapped against the mouse reference genome (build 37) using TopHat2^45^. The aligned reads were assembled into transcripts, and gene expression was calculated using Cufflinks^46^. The expression level of genes was determined based on the value of fragments per kilobase per million (FPKM), which was calculated as the number of fragments mapped to the transcripts of one gene divided by the transcript length and the number of total mapped reads in one sample. The differentially expressed genes were identified using Cufflinks (Fold change>4, FDR<0.01)^46^. Principal component analysis was carried out with the web tool ClustVis^47^.

Human orbital fat (n=2) was obtained from patients undergoing routine resection of prolapsed orbital fat, which has been shown to be intraconal fat^48^, using a protocol approved by the Johns Hopkins University Institutional Review Board and following the tenets of the Declaration of Helsinki. RNA was extracted from each fat depot using Trizol and RNeasy mini kits (Qiagen, Hilden, Germany). RNA-Sequencing libraries were made using stranded Total RNASeq library prep. Libraries were sequenced with Illumina NextSeq500, paired-end read of 75 bp, 50 million reads per library. A bioanalyzer was used to perform quality control. cDNA libraries were prepared for samples with RIN>6 and RNA sequencing performed using Illumina Nextseq500. Illumina adapters of libraries after sequencing were removed using Cutadapt (v1.18;^49^) with default parameters. Libraries were then aligned to GRCh38 using STAR (v2.42a;^50^) with –twopassMode Basic. RSEM (v1.3.1;^51^) was used for transcript quantification, with rsem-calculate-expression (--forwad-prob 0.5). TPM value from RSEM was used to plot white, beige, or brown adipocytes-enriched markers using ggplot (v3.32).

### Thermoneutral adaptation

Six-week old CD-1 and C57BL/6 males were housed at 30°C for 4 weeks. The mice were then euthanized, and the orbital soft tissues were dissected and fixed in 4% PFA. The tissue was embedded in paraffin and sectioned. Haematoxylin and eosin (H&E) staining of paraffin-embedded sections was used to visualize orbital fat, as described^52^.

### Comparative Anatomy

Orbital fat from cat and dog eyes, which were obtained following methods in full accordance with protocols approved by the Michigan State Animal Care and Use Committee, were provided by Simon Petersen-Jones (Michigan State University School of Veterinary Medicine), and fixed in 4% PFA following dissection. Male Golden Syrian hamsters housed at room temperature were obtained from Envigo and perfused with 4% PFA at 9-10 weeks of age. Orbital fat was dissected and fixed in PFA overnight. Lamb and goat orbital fat was dissected from animal heads obtained from International Grocery and Halal Meat INC (Baltimore, MD). Human orbital fat specimens submitted were obtained from patients undergoing orbital or peri-ocular surgery under a protocol approved by the Johns Hopkins University Institutional Review Board following the tenets of the Declaration of Helsinki, with informed consent obtained for all samples analyzed. H&E staining of paraffin-embedded sections was used to visualize orbital fat^52^.

### Microscopy

Imaging was performed using a Keyence BZ-X710 microscope (Keyence, Osaka, Japan).

### Quantitative RT-PCR

Whole RNA was extracted from mouse orbital fat samples using RNeasy mini-kits (Qiagen). cDNA was prepared using Superscript III first-strand synthesis kit (ThermoFisher). Quantitative PCR was performed in triplicate using on a … using the following primer sets: *PPIA* (CAGACGCCACTGTCGCTTT, TGTCTTTGGAACTTTGTCTGCAA), *Ucp1* (AGGCTTCCAGTACCATTAGGT, CTGAGTGAGGCAAAGCTGATTT), *Cidea* (TGACATTCATGGGATTGCAGAC, GGCCAGTTGTGATGACTAAGAC), *Gdf5* (CCCATCACACCCCACGAATA, CTCGGTCATCTTGCCCTTTG). *PPIA* was chosen as a reference gene after comparing the expression of *TBP*, *PPIA*, and *GAPDH* in brown fat, inguinal fat, epididymal fat, and orbital fat using RefFinder^53^ (https://www.heartcure.com.au/reffinder/). Data were compared using the delta-delta method. One-way ordinary ANOVA of data was performed using Prism (GraphPad).

### *In situ* hybridization

Single-molecule *in situ* hybridization was performed using the following RNAscope probes: Mm-Ucp1 (catalog # 455411), Mm-Gdf5 (catalog # 407211), Mm-Cxcl12-C2 (catalog # 422711-C2) (Advanced Cell Diagnostics, Newark, CA) as per manufacturer’s protocol.

## Data availability

All sequencing data are available on GEO (GSE158464).

## Acknowledgements

We would like to thank Subhash Kulkarni for sharing the Wnt1:Cre mouse line. We would also like to thank Susan Aja and Michael Wolfgang for assistance with housing mice at thermoneutral temperature, and Transcriptomics and Deep Sequencing Core (Johns Hopkins) for the preparation and sequencing of RNA-Seq libraries. FR is supported by the National Eye Institute of the National Institutes of Health under award number K08EY027093. SB is supported by NIH Grant R01EY020560. DWK is supported by the Maryland Stem Cell Research Fund (2019-MSCRFF-5124).

## Author Contributions

F.R. and S.B. conceived and oversaw the study. F.R. conducted RNA-Seq, qRT-PCR and histological analysis of orbital fat. D.W.K. analyzed RNA-Seq data and performed single-molecule fluorescent *in situ* hybridization. J.P. and J.Q. analyzed RNA-Seq data. N.R.M. and C.G.E. provided human orbital fat samples. F.R. and S.B. wrote the manuscript, with input from all authors.

---

**Figure S1. RNA-Seq analysis of extraocular muscles identifies contaminating transcripts in orbital fat samples. A.** Principal component analysis of transcriptome data from mouse brown fat (BF), epididymal fat (EF), inguinal fat (IF), orbital fat (OF) and extraocular muscle (EOM) demonstrates that BF and OF segregate more closely to each other than to EF and IF. EOM segregates separately from all fat depts. **B.** Sample-wise correlation of transcriptome data from BF, EF, IF and OF demonstrates that transcriptome of OF is more closely correlated with that of BF than with the transcriptomes of EF, IF, and EOM. **C.** Heat map of transcriptome data from mouse BF, EF, IF, OF, and EOM demonstrating that there is little overlap between EOM and OF.

**Table S1: List of genes selectively enriched in retrobulbar fat.**

