## Supplementary material for "Temperature and species-dependent regulation of browning in retrobulbar fat": Figure S1 and Table S1

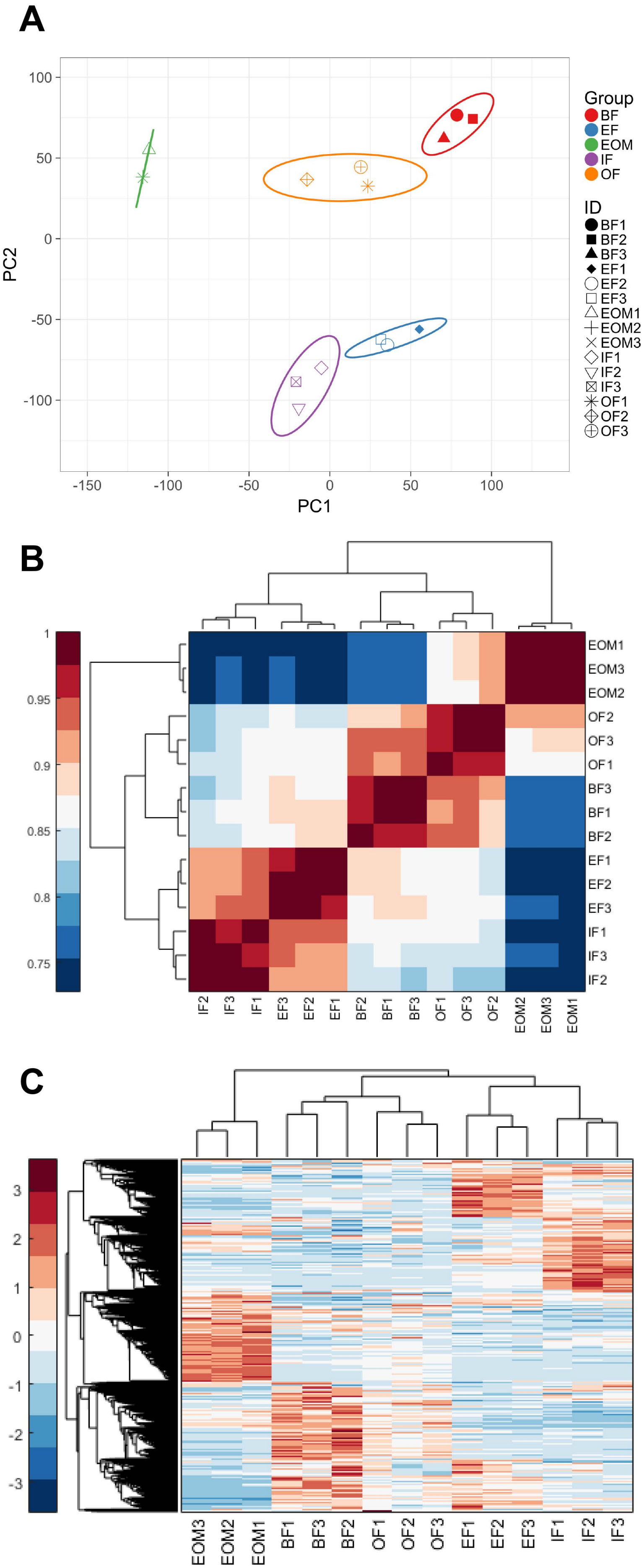


Figure S1 Rajaii, et al.

| Gene ID | Mean OF | Mean BF | Mean EF | Mean IF | Mean EOM | p-value | FDR | log2 Fold Change |
|---|---|---|---|---|---|---|---|---|
| *Gldn* | 3.12 | 2.15 | 2.64 | 2.64 | 0.64 | 3.0E-05 | 0.04 | 2.12 |
| *Actn3* | 90.57 | 3.39 | 46.98 | 46.98 | 83.96 | 3.6E-04 | 0.09 | 2.02 |
| *Serpini1* | 2.00 | 1.20 | 1.60 | 1.60 | 1.03 | 1.7E-06 | 0.02 | 1.50 |
| *Cxcl12* | 72.52 | 15.18 | 43.85 | 43.85 | 60.00 | 1.2E-04 | 0.07 | 1.32 |
| *1200009I06Rik* | 1.60 | 0.17 | 0.89 | 0.89 | 1.42 | 4.0E-04 | 0.09 | 1.31 |
| *Gm10193* | 2.57 | 1.56 | 2.07 | 2.07 | 1.17 | 1.3E-04 | 0.07 | 1.24 |
| *Slc16a2* | 11.32 | 7.45 | 9.39 | 9.39 | 6.67 | 4.6E-05 | 0.05 | 1.24 |
| *Prickle4* | 0.98 | 0.80 | 0.89 | 0.89 | 0.43 | 3.7E-04 | 0.09 | 1.17 |
| *Fst* | 17.57 | 4.80 | 11.19 | 11.19 | 14.41 | 4.6E-04 | 0.09 | 1.04 |
| *Plscr4* | 2.04 | 2.55 | 2.30 | 2.30 | 4.56 | 3.4E-05 | 0.04 | -1.04 |
| *Nedd9* | 3.89 | 4.57 | 4.23 | 4.23 | 7.42 | 4.2E-04 | 0.09 | -1.14 |
| *Plk3* | 0.60 | 1.06 | 0.83 | 0.83 | 0.98 | 1.5E-04 | 0.07 | -1.16 |
| *Hip1* | 4.48 | 5.15 | 4.81 | 4.81 | 8.17 | 4.2E-04 | 0.09 | -1.21 |
| *Ptger4* | 1.02 | 1.96 | 1.49 | 1.49 | 1.80 | 4.6E-04 | 0.09 | -1.23 |
| *Pde7a* | 3.51 | 3.66 | 3.59 | 3.59 | 10.21 | 2.2E-04 | 0.08 | -1.29 |
| *Plau* | 7.31 | 8.08 | 7.69 | 7.69 | 25.57 | 3.9E-04 | 0.09 | -1.51 |
| *Nap1l5* | 0.51 | 2.51 | 1.51 | 1.51 | 1.61 | 1.4E-04 | 0.07 | -1.73 |
| *Spon2* | 0.89 | 1.15 | 1.02 | 1.02 | 3.82 | 4.5E-04 | 0.09 | -2.02 |
| *Galnt12* | 0.20 | 0.26 | 0.23 | 0.23 | 0.93 | 2.6E-04 | 0.08 | -2.05 |
| *Hoxd4* | 0.19 | 0.78 | 0.48 | 0.48 | 0.48 | 1.7E-04 | 0.07 | -2.10 |
| *Hoxd8* | 0.73 | 6.62 | 3.68 | 3.68 | 2.03 | 1.1E-04 | 0.07 | -2.58 |
| *Meis2* | 0.21 | 2.44 | 1.33 | 1.33 | 0.24 | 1.3E-04 | 0.07 | -3.26 |
| *Hoxb3* | 0.02 | 1.37 | 0.69 | 0.69 | 0.05 | 2.6E-04 | 0.08 | -5.32 |
| *Hoxb4* | 0.02 | 2.51 | 1.26 | 1.26 | 0.03 | 2.9E-04 | 0.08 | -6.10 |
| *Hoxb2* | 0.01 | 2.06 | 1.04 | 1.04 | 0.01 | 2.0E-04 | 0.07 | -7.65 |
| *2700033N17Rik* | 0.00 | 0.95 | 0.48 | 0.48 | 0.12 | 6.1E-05 | 0.05 | |

Table S1 Rajaii, et al.
